# The long-distance flight behavior of *Drosophila* suggests a general model for wind-assisted dispersal in insects

**DOI:** 10.1101/2020.06.10.145169

**Authors:** Katherine Leitch, Francesca Ponce, Floris van Breugel, Michael H. Dickinson

## Abstract

Despite the ecological importance of long-distance dispersal in insects, its underlying mechanistic basis is poorly understood. One critical question is how insects interact with the wind to increase their travel distance as they disperse. To gain insight into dispersal using a species amenable to further investigation using genetic tools, we conducted release-and-recapture experiments in the Mojave Desert using the fruit fly, *Drosophila melanogaster*. We deployed chemically-baited traps in a 1 km-radius ring around the release site, equipped with machine vision systems that captured the arrival times of flies as they landed. In each experiment, we released between 30,000 and 200,000 flies. By repeating the experiments under a variety of conditions, we were able to quantify the influence of wind on flies’ dispersal behavior. Our results confirm that even tiny fruit flies could disperse ∼15 km in a single flight in still air, and might travel many times that distance in a moderate wind. The dispersal behavior of the flies is well explained by a model in which animals maintain a fixed body orientation relative to celestial cues, actively regulate groundspeed along their body axis, and allow the wind to advect them sideways. The model accounts for the observation that flies actively fan out in all directions in still air, but are increasingly advected downwind as winds intensify. In contrast, our field data do not support a Lévy flight model of dispersal, despite the fact that our experimental conditions almost perfectly match the core assumptions of that theory.

**Significance Statement:** Flying insects play a vital role in terrestrial ecosystems, and their decline over the past few decades has been implicated in a collapse of many species that depend upon them for food. By dispersing over large distances, insects transport biomass from one region to another and thus their flight behavior influences ecology on a global scale. Our experiments provide key insight into the dispersal behavior of insects, and suggest that these animals employ a single algorithm that is functionally robust in both still air and under windy conditions. Our results will make it easier to study the ecologically important phenomenon of long-distance dispersal in a genetic model organism, facilitating the identification of cellular and genetic mechanisms.

## Introduction

If asked to picture a migrating insect, the first image that comes to mind might be a large charismatic species such as the monarch butterfly, whose seasonal movements across North America have inspired naturalists for centuries. However, as pointed out by David and Elizabeth Lack(1), our impression of insect migration is strongly biased toward large animals; many species are so small that their geographic relocations escape our attention, especially if their population densities are not strongly concentrated by geological features such as narrow mountain passes. As research using high-altitude traps(2) and upward looking radar(3) indicates, long distance migration may be more ubiquitous and ecologically important among both large and small insects than previously appreciated. Long-distance dispersal - i.e. the non-cyclic movement from one area to another, is even harder to observe and study in small insects, because the events are not generally predictable and the animals are far too small to be captured on radar or outfitted with tracking devices. Understanding long-distance migration and dispersal is quite important, however, because these phenomena provide an important means by which biomass relocates on both local and global scales(4). Furthermore, as insect population densities decline due environmental degradation and climate change(5–7), understanding the dispersal capacity of insects and the behavioral algorithms that underlie them will be crucial in predicting the ecological impact of population decline.

Although not generally renowned for its capability to disperse over long distances, a series of release-and-recapture experiments over 40 years ago suggest that the fruit fly, *Drosophila melanogaster*, may be capable of movements on the order of 15 kilometers in a single night, a distance equivalent to 6 million body lengths(8, 9). These experiments were conducted by releasing tens of thousands of fluorescently labeled flies in the evening, and then censusing the contents of traps baited with yeast and banana placed at distant oases the next morning. While these pioneering studies suggested that the dispersal capacity of *Drosophila* was much greater than previously estimated, they left open several critical questions. First, it was not clear whether individual flies dispersed in random directions, or whether the population movement was biased by external conditions, such as the wind, geographical features, or celestial cues. Second, because the precise transit times of the flies were not known, it was impossible to estimate the actual groundspeeds used by the animals as they dispersed. To provide more clarity to these and other questions related to long-distance dispersal, we conducted a series of release-and-recapture experiments in the Mojave Desert. We equipped circular arrays of chemically-baited traps with simple machine vision systems that captured the arrival times of flies as they landed, and repeated the experiments under a variety of ambient wind conditions. The results provide key insight into the behavioral algorithms used by *Drosophila* while dispersing in the wild, and serve as the basis for a general model of wind-assisted dispersal in insects.

## Results

To examine the long-distance flight behavior of *Drosophila*, we performed a series of release-and-recapture experiments on a dry lakebed, Coyote Lake, in the Mojave Desert. We situated the release site near the center of the lakebed, and deployed a circular ring of baited traps, each equipped with a camera aimed at the trap surface (Fig 1A). The bait consisted of a fermenting solution of sugar and apple juice that actively produced CO_2_ and ethanol, which we suspect served as the primary long-distance attractants in our experiments(10). In an initial trial, we positioned eight traps at 250 m from the release site; but in all subsequent experiments (N=5), we positioned ten traps at a radius of 1 km. The 28 × 28 cm mesh surface of the traps contained an array of inwardly pointed mesh funnels that allowed entry, but limited egress, thus allowing us to count and identify the flies at the end of the experiment. (Fig 1B, C). An anemometer placed at the release site recorded instantaneous windspeed and direction over the course of each experiment. In preliminary releases to test our protocol, we marked the flies with fluorescent dust as had been done in previous studies(8, 9). However, after we found no evidence that *D. melanogaster* were ever present at the lake bed (unless we released them), we did not mark the flies as the presence of the fluorescent power could interfere with the animals’ behavior and flight performance. The number of flies released in each experiment ranged from ∼30,000 to 200,000, consisting of both males and females.

**Figure 1.**
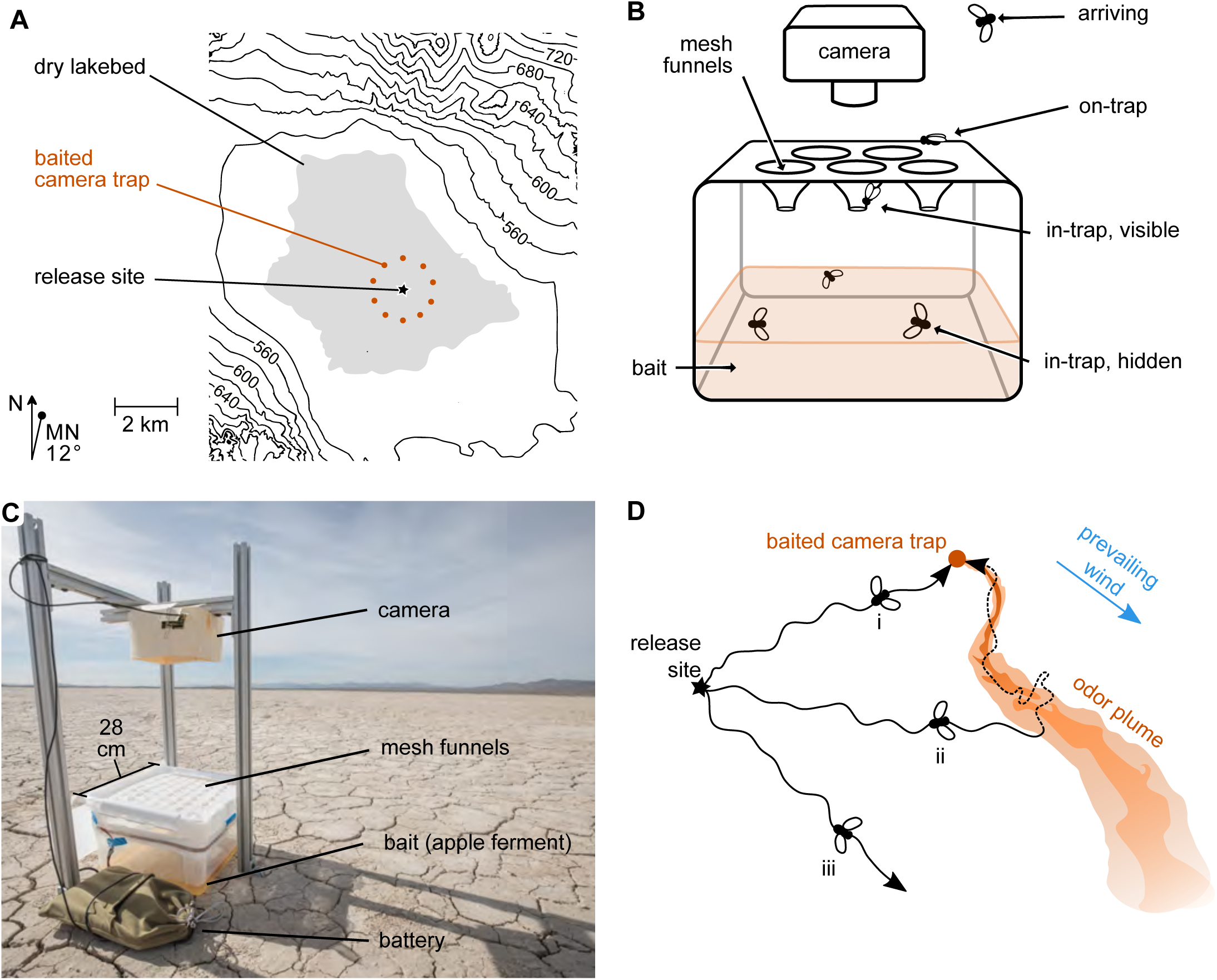
Experimental protocol for field release and monitored recapture of laboratory-reared *Drosophila*. **A**. We conducted field experiments at Coyote Lake (gray area), a dry lake bed in the Mojave Desert of southern California. The contour values are given in meters; the lake bed surface is ∼520 m above sea level. Our field paradigm typically used ten baited camera traps (orange) at a radius of 1 km from a central release site (black star). North, N; magnetic north; MN. **B**. Cartoon of odor-baited camera trap. The top surface consists of a polyester mesh plane with an array of mesh funnels projecting inward toward the apple-yeast ferment (bait). For simplicity, only 5 funnels are drawn, the actual traps contained 60 funnels. The camera mounted above captures time-lapse images of the trap, measuring the number of flies atop the trap (on-trap) and the number in a zone directly on the underside of the mesh (in-trap, visible). The camera cannot detect flies that are deeper inside the trap (in-trap, hidden) or those arriving at the trap before they land (arriving). However, all flies in the trap may be counted at the end of the experiment. Dimensions not to scale. **C**. Photograph showing one odor-baited camera trap deployed on the lakebed. The desiccation polygons visible here are representative of the lakebed surface. The southeastern terminus of the Calico Mountains, roughly 7 km away, is visible on the horizon. **D**. A top-view cartoon illustrating key assumptions guiding our experimental design. Wind (blue arrow) advects a turbulent plume of attractive odor from each baited camera trap (one trap shown, orange). We assume that the ability of a fly to detect and track a plume falls off with distance from the source (illustrated by orange gradient). Flies disperse from the release site (black star), each maintaining a particular trajectory of some mean direction and some tortuosity (black lines). Flies whose trajectories happen to intersect the plume near a trap (i) are likely to be among the earliest arrivers, owing to their relatively direct flight path. Flies that intercept a plume downwind of its source (ii) will follow a longer flight path before arriving at the trap, given the distance added by plume tracking (broken line). Many flies (iii) may never encounter a detectable plume.

### Flies disperse randomly in low wind, but have a downwind bias in the presence of wind

We suspect that a small subset of flies would adopt trajectories leading directly to a trap (Fig 1D, i), but that the majority of those recaptured would first encounter a trap’s odor plume, and then track it upwind to find the source (Fig 1D, ii). Thus, we assume the trap count distributions to reflect two processes: 1) a dispersal flight over open space before a fly detects an odor plume, and 2) an upwind flight within the plume toward the trap, presumably mediated by the well characterized cast-and-surge behavior exhibited by *Drosophila* and other animals (11). Wind is expected to impact both these processes by shaping the odor plumes and by influencing the animal’s groundspeed. Fortuitously, the wind conditions were different on every day we conducted a release, allowing us to examine the influence of windspeed by comparing the results from different experiments. We found that mean windspeed influenced the final trap count distributions in a systematic manner (Fig 2A, B). Gentle winds resulted in a more uniform distribution of trap counts around the ring (Fig. 2C), while at higher average windspeeds, the distributions were skewed in a downwind direction. Despite this downwind bias in trapping distribution, some flies always managed to arrive at crosswind and upwind traps in all but the strongest of wind conditions. Although we could only roughly estimate the total number of flies we released, we note that higher windspeeds reduced the percentage of released flies that we recaptured at the 1-km traps, which varied from 1.3 to 0.1% (Fig. 2A). On one particularly windy day (> 5 m s^-1^), we only recovered 0.01% of the flies (data not shown), which did not yield enough data to analyze. Collectively, these results indicate that *Drosophila*, despite their small size, are not simply advected by the wind as they disperse, but rather have some capacity to fly in upwind and crosswind directions. We did not happen to conduct releases on any day when the windspeed was zero, but by extrapolating to this condition, we conclude that in the absence of wind flies tend to fan out randomly in all directions when released (Fig. 2C).

**Figure 2.**
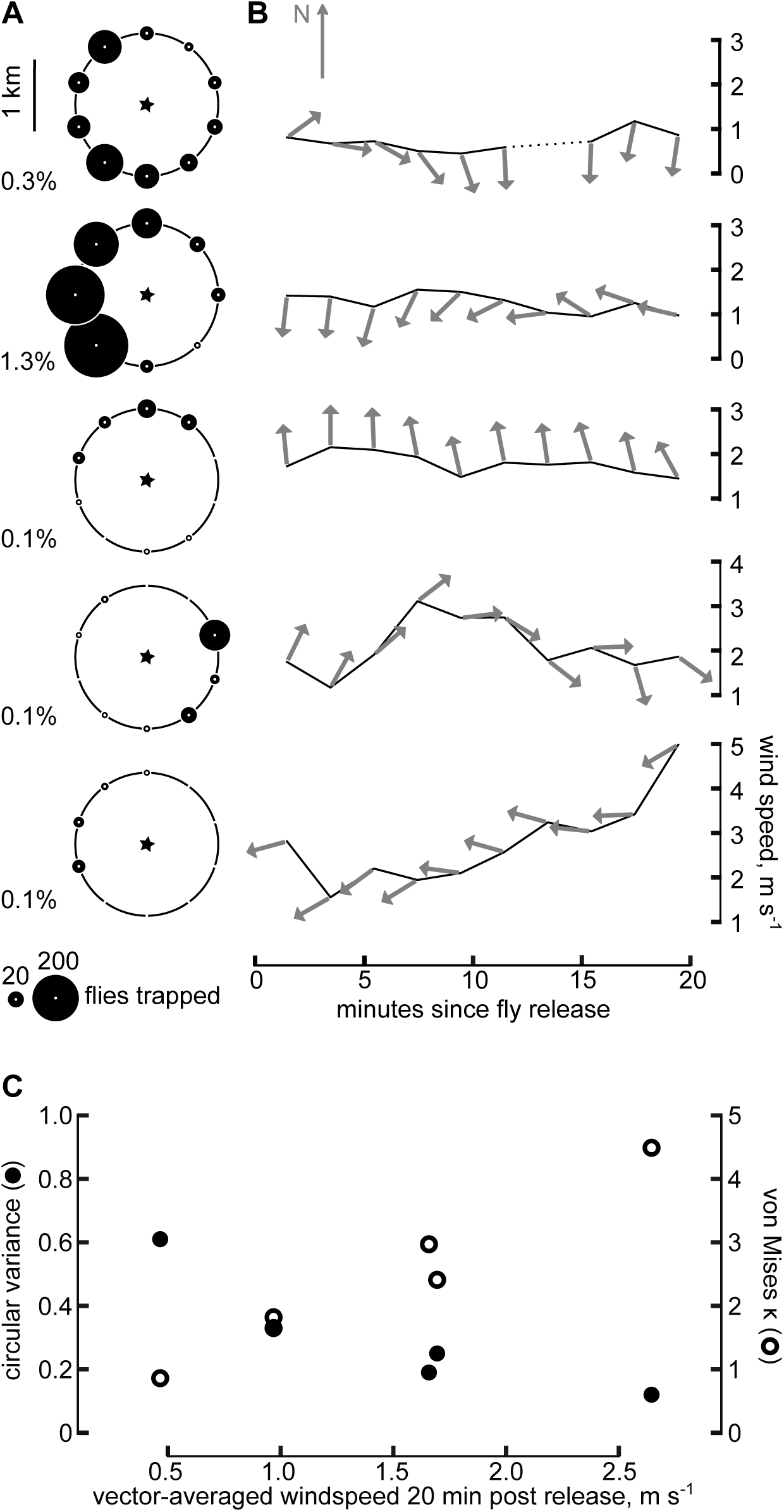
Influence of wind on circular distribution of trapped flies. **A**. Birds-eye representations of trap-count data, with geographic north oriented up. Traps were positioned 1 km away from the release site (black star). Each row depicts the results of a different field release. The area of each black circle indicates the number of flies caught at that trap by the end of each experiment. The five experiments depicted here are ordered according to the windspeed, vector-averaged for the first 20 minutes post-release (top row, lowest windspeed). Recapture percentages, calculated using estimates of release populations, are indicated at bottom left of each panel. **B**. Windspeed and direction during each of the field releases depicted in (A). Data were vector-averaged in 2-minute bins. Arrows point in the downwind direction; geographic north is oriented up. For clarity, arrow lengths are fixed; the position along the ordinate of each arrow’s base denotes vector-averaged windspeed. We recorded wind data continuously, barring one case of anemometer failure (first row, gap in data from 10 to 14 minutes, post-release). **C**. Two measures of distribution statistics, circular variance (closed circles) and the von Mises parameter Κ (open circles), are plotted as a function of vector-averaged windspeed for the five experiments shown above.

### Flies maintain a constant groundspeed, whether going up or downwind

To determine if *Drosophila* actively regulate their groundspeed during dispersal, we compared arrival dynamics at downwind and upwind traps as measured by the cameras. We pooled the camera data collected from the circular array of traps into two groups, representing the downwind and upwind sectors (Fig 3, black vs. green data). This analysis was conducted on the data collected using an array of traps 250 m from the release site, because the higher proportion of flies recaptured at this shorter distance (∼2%) provided more arrival events. Although the pooled data from upwind and downwind traps differed with respect to the total number of flies captured, the time courses of arrivals were remarkably similar. This similarity was apparent not only in the on-trap data (Fig 3A), which reflect the rate of arrival, but also in the in-trap data (Fig 3B), which approximate the cumulative number of flies having arrived at the traps. These findings suggest that the recaptured flies, despite having taken widely different trajectories relative to the prevailing wind of ∼1.5 m s^-1^, had managed to achieve roughly similar groundspeeds. This ability of flies to regulate flight speed relative to the ground in the face of varying winds is well known from wind tunnel experiments(12). Flies adjust groundspeed primarily by adjusting body pitch(13, 14), which they regulate via feedback from translational optic flow(15, 16).

**Figure 3.**
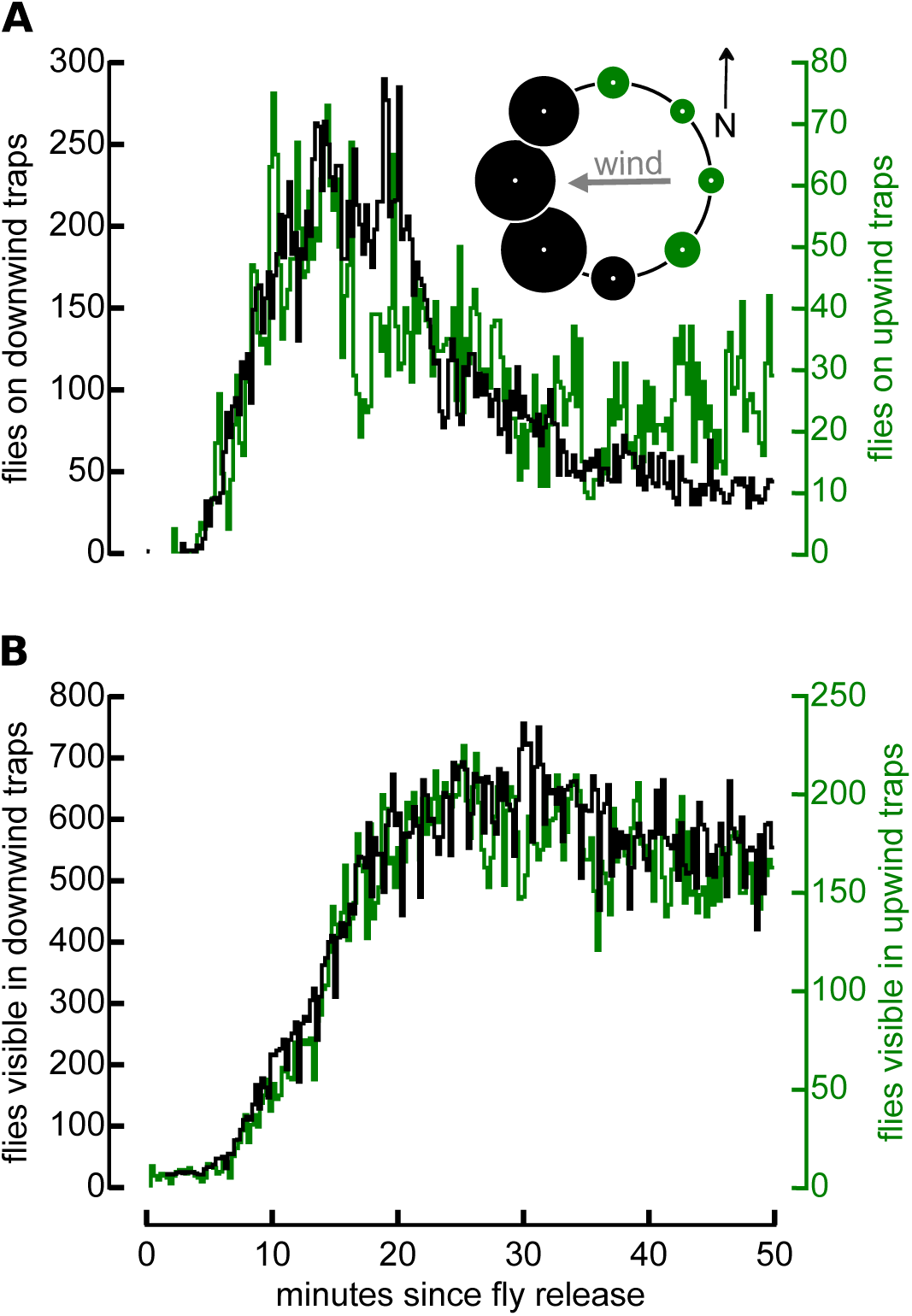
In moderate wind, *Drosophila* arrive at upwind and downwind traps with a similar time course. **A**. During this experiment, the traps were positioned at a 250 m radius from the release site instead of 1 km. The vector-averaged wind over 20 minutes following the fly release was 1.5 m s^-1^, from slightly north of due east (gray arrow, inset at right). Based on this mean direction, we define four traps as upwind (green circles, inset) and the remaining four as downwind (black circles, inset). The number of flies imaged on the trap surface (black line, left axis), summed across all downwind traps for each time bin, is shown with the number of flies imaged on top of all upwind traps (green line, right axis; note difference in scale). **B**. From the same experiment, we also imaged the number of flies visible within the upwind (green line, right axis) and downwind (black line, left axis) traps; these time-series data are concurrent with the data in (A). Note that this trap-count distribution appears somewhat similar to that in Fig. 2A, row 2, but these data come from different experiments, on different days.

### What is the groundspeed flies actually use?

The small distance between release site and trap in the 250 m experiment made it hard to accurately estimate the magnitude of flies’ groundspeeds, because the flight time was so brief. By analyzing the data from experiments using traps set at 1 km, we could derive a better estimate of the flies’ groundspeed, as well as examine the influence of the wind more accurately. Our strategy was to estimate the groundspeed of the first flies to arrive at each trap (hereafter, ‘first arrivers’), as these were individuals that most likely flew directly to the trap without requiring an extended bout of upwind plume tracking. Unfortunately, we could not use our automated machine vision analysis to determine the flight time of the first arrivers in most cases, because changes in lighting and movements of the mesh surface caused by the wind generated false positives that compromised our ability to score rare events. This was particularly problematic when relatively few flies came to a trap. For this reason, we manually annotated the camera images to determine the first appearance of a *Drosophila-*shaped insect. Thirty different traps from five experiments were amenable to this analysis, providing a measure of groundspeed as a function of windspeed (Fig. 4C). The camera data from 9 of these 30 traps provided a sufficiently high signal-to-noise ratio that we could compare the manually annotated arrival time with an automated analysis and the correspondence was reasonably good (Supplemental Figure 1).

**Figure 4.**
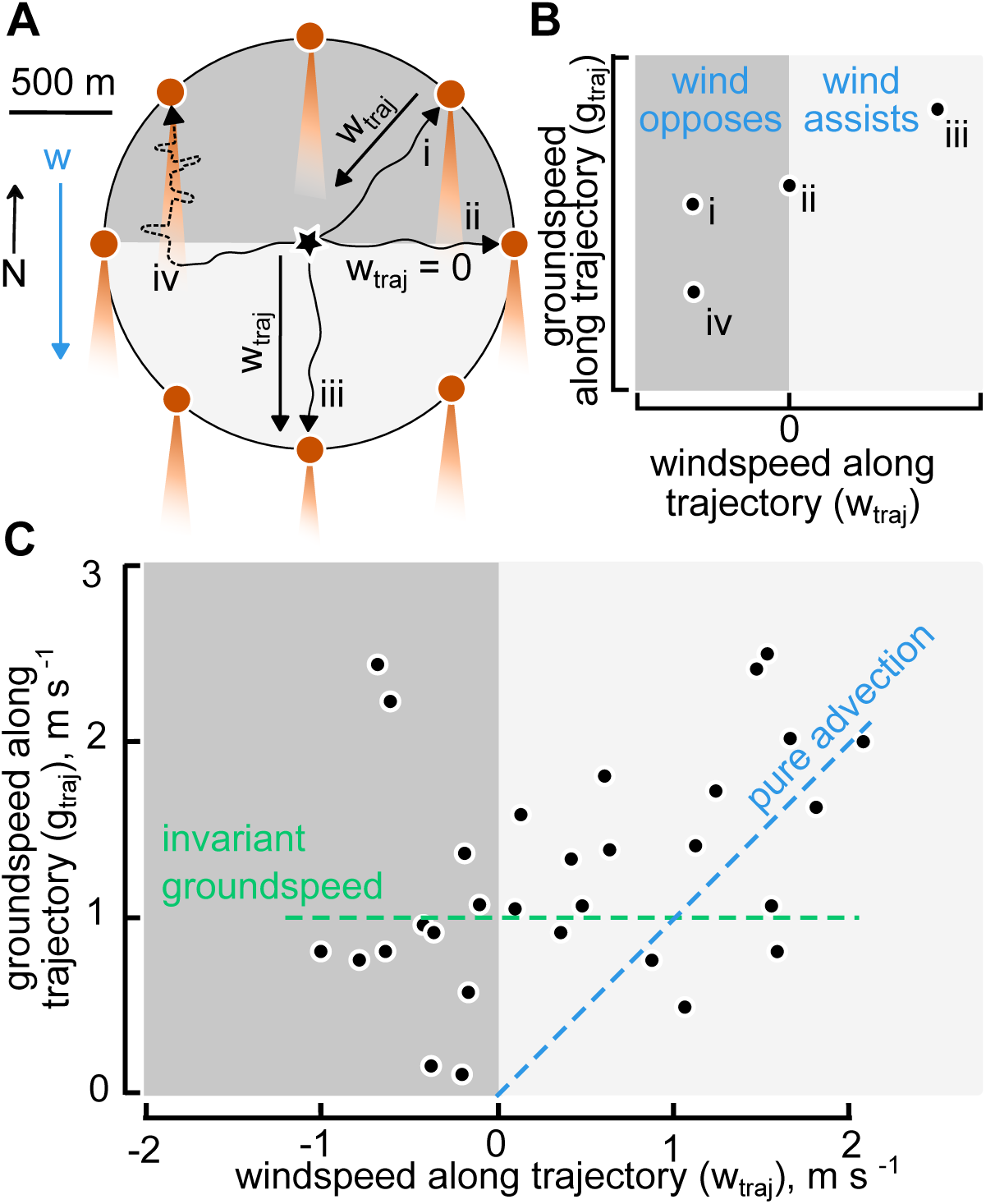
Our method to examine the influence of wind on groundspeed. **A**. Hypothetical trajectories (meandering black lines) of four flies leaving the release site (star). Wind (w, blue vector) advects odor plumes (orange wedges) due south of each trap (orange circles). We calculated flies’ groundspeeds using each measured arrival time and average trajectory, with the latter approximated as the straight path from the release site to the trap. This heuristic is most appropriate for flies whose trajectories happen to intercept a plume very close to the trap (flies i-iii). The farther downwind a fly encounters the odor plume, the less accurate this assumption becomes (e.g. fly iv). Black vectors show the component of wind parallel to trajectories i - iii (w_traj_). **B**. For each fly in (A), we plot the groundspeed along its average trajectory (g_traj_) as a function of windspeed along this average trajectory (w_traj_). Negative w_traj_ values (dark gray) denote trajectories opposed by the wind; positive (light gray) indicate trajectories assisted by prevailing wind. g_traj_ values reflect a model in which flies attempt to regulate their groundspeeds over a range of wind conditions (flies i and ii), although at higher values of w_traj_ advection likely dominates (fly iii). g_traj_ values will be underestimated for flight paths that do not intercept a plume near a trap (fly iv). **C**. Field measurements (black points) of g_traj_ and w_traj_, as described above, from 30 traps over five experiments (same experiments as in Fig. 2). Superimposed are the relationships expected if flies were entirely advected by wind (blue) or if flies perfectly maintained some preferred groundspeed along their trajectory (green).

From our measurements of arrival time, we determined the groundspeed of the first arrivers (*g*_*traj*_), assuming that these flies took off immediately upon release and flew in a straight line to the trap. Across all our experiments and traps, these first arrivers flew with an average groundspeed of 1.4 ± 0.6 m s^-1^, however, the precise values were also a function of the wind.

### The influence wind on groundspeed

From our anemometer data, we determined the average headwind or tailwind that the first arrivers would have experienced along their straight-line trajectories, *w*_*traj*_. Plotting the first arrivers’ average ground speeds against the wind speed along their trajectories (Fig. 4B), indicates the influence of the wind conditions. If the flies were simply advected by the wind, all the data should lie on a line running through the origin, with a slope of one. If the flies compensated groundspeed perfectly, then the data should fall about a horizontal line intersecting the ordinate axis at some preferred groundspeed value. The distribution of field data suggests an intermediate behavior, that is, flies exhibit some ability to actively regulate groundspeed at low windspeeds, but tend to move with the wind as windspeed increases. Two data points (within the range of negative *w*_*traj*_ values) showed unusually large values for groundspeed, and we suspect that these outliers were due to a misidentification of some local fly-sized insects when we annotated the trap images. All subsequent analyses were performed on both the complete thirty-point data set and after excluding these two outliers, but the exclusion did not alter any qualitative conclusions of our study.

The cluster of groundspeed values measured when the windspeed along the trajectory was near zero provides a rough estimate of the preferred groundspeed that flies use in the absence of any wind. This value, ∼1 m s^-1^, is much higher than measurements of *Drosophila* flight velocities made in indoor wind tunnels(17), but consistent with values made for flies flying in green houses(18). Obviously, any first arrivers that lingered at the release site, flew in a zig-zag manner, or had to work upwind long distances within the odor plume to reach the trap would have to have flown even faster to arrive at the recorded time (Fig. 4A). Thus, our calculations are estimates for the absolute minimum flight speed the animals could have used. The relatively high value suggests that the flies – the first arrivers at least – must have flown in rather straight trajectories between the release site and traps. If they had executed highly meandering flight paths – those predicted by a random search pattern, for example - they could not have reached the traps at the recorded times unless they had flown at groundspeeds that were far in excess of what is physically possible.

### Wind-assisted dispersal model

Thus far, our results suggest that a population of flies fans out in different directions when released, with at least some flies maintaining a roughly straight trajectory. We suggest two basic mechanisms by which the flies might maintain a straight path as they disperse. In the first, each fly chooses a constant heading (i.e. body orientation) relative to a celestial cue. In the second, flies somehow actively maintain a constant trajectory (i.e. a straight path over the ground), perhaps by orienting to a distant visual landmark. Note that these two hypotheses are distinct because a flying animal’s trajectory need not be aligned to its heading, i.e. it can move such that its longitudinal body axis (heading) is not aligned with its groundspeed vector (trajectory). The constant trajectory strategy has been observed in bumblebees, which actively compensate for wind drift during their nest-bound flights(19), and partial compensation has been observed in migratory noctuid moths(20). The constant heading hypothesis is consistent with radar observations of hoverflies’ autumn migrations(21), and with laboratory experiments showing that tethered flies will maintain a fixed orientation relative to patterns of polarized light(22–24) or a small bright spot simulating the sun(25). Furthermore, these lab experiments indicate that each fly appears to choose an arbitrary heading from random that it then maintains over time, which could easily explain why the flies fanned out in different directions upon release.

Celestial menotaxis alone cannot explain why flies are biased downwind as the wind increases in strength. To account for the influence of the wind, we propose a simple behavioral algorithm that could explain the key features of our field data and is consistent with prior laboratory observations (Fig. 5A). In our model, each fly maintains a constant heading and regulates groundspeed as our data suggest, but the groundspeed regulator only operates on the velocity component oriented along the body axis. In other words, the fly does not regulate sideslip, but rather allows itself to be advected sideways due to the wind. We developed a set of four models that all incorporated unregulated sideslip, but with different variations, thus creating a 2 × 2 matrix of two binary assumptions: 1) fixed random heading vs. fixed random trajectory, and 2) regulated groundspeed along the body axis vs. unregulated groundspeed (Fig. 5). Figure 6A–C shows how three example flies differing only in their chosen heading angle would behave according to model A (fixed heading with regulated groundspeed, Fig. 5A), and how these would relate to the values of *w*_*traj*_ and *g*_*traj*_ available from field measurements (see sample points in Fig. 4B). These illustrations are equivalent to three simulations of model A; running this simulation over a wide range of windspeeds and directions generates a full set of predictions. The contours of the simulations are constrained by three free parameters: the minimum (air_min_) and maximum (air_max_) airspeed achievable by the flies and the preferred groundspeed (g_pref_) (Fig. 6D); these values were drawn from velocity statistics of *Drosophila* in free-flight, in brightly lit greenhouses(18). Figure 6D shows the output of model A overlaid with our field data (n = 30, from 5 different releases). The other three models implemented the other pairs of assumptions: model B, unregulated groundspeed with fixed heading; model C, regulated groundspeed with fixed trajectory; model D, unregulated groundspeed with fixed trajectory. In all four models, sideslip was unregulated.

**Figure 5.**
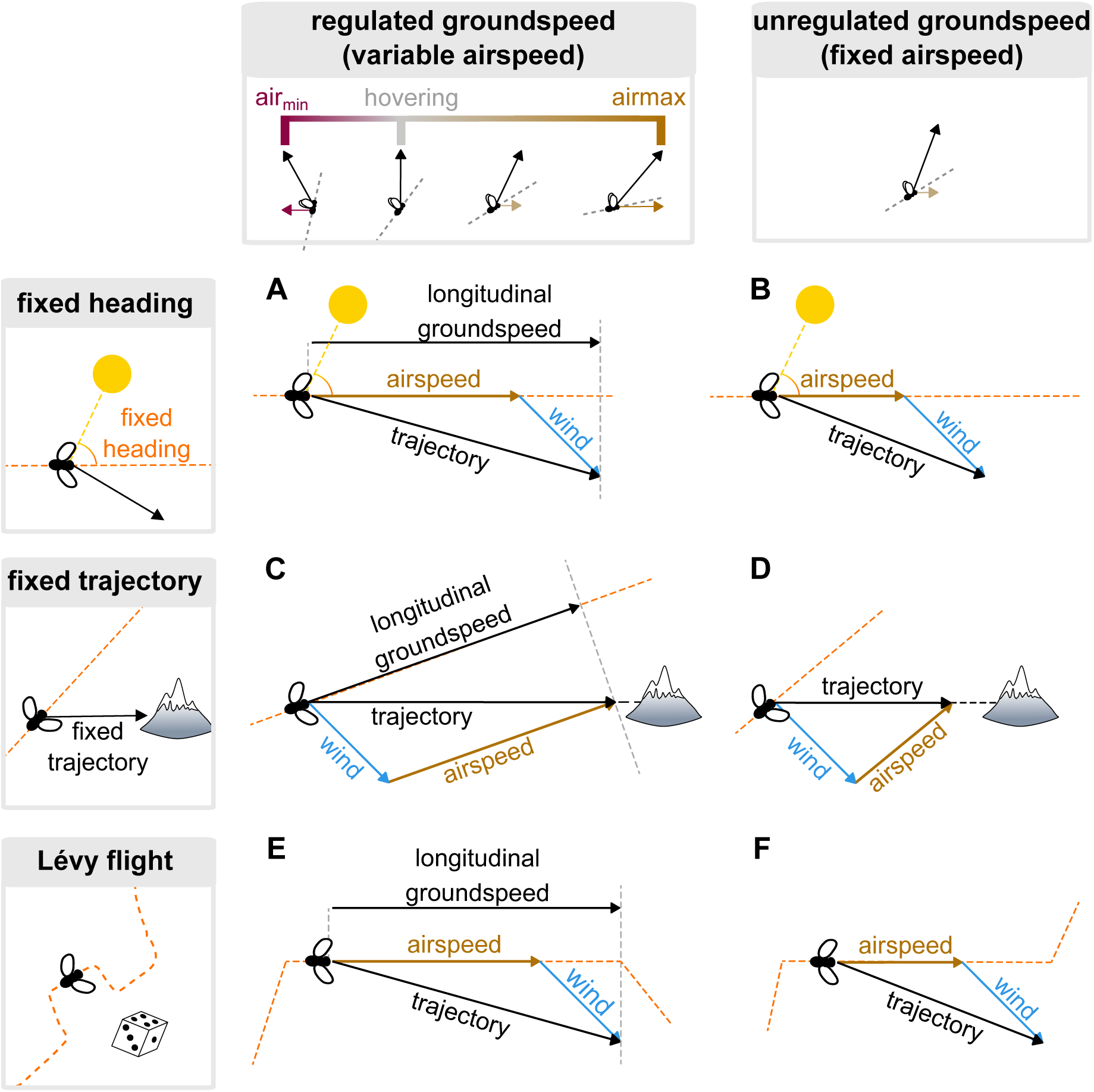
Six behavioral models of free-flight navigation in the presence of wind. **A – F**. Cartoons depicting how a fly’s heading (broken orange lines), trajectory, longitudinal groundspeed, and airspeed relate to the wind in each of six behavioral models. These simple models differ in azimuthal orientation strategies (rows) and in the presence or absence of groundspeed regulation (columns). **A**. Model A, in which each fly maintains a fixed body angle (“fixed heading”, top row) relative to an external azimuthal reference (depicted as a sun), and adjusts its airspeed, within limits, to achieve a preferred longitudinal groundspeed (“regulated groundspeed”, left column). The fly’s airspeed along its body axis (brown vector) sums with the total wind vector (blue), generating the fly’s trajectory (black). Projecting the trajectory vector onto the fly’s body axis gives the fly’s longitudinal groundspeed (top black vector), which the fly actively regulates. **B**. Model B, in which each fly maintains a fixed heading as in the previous model, but does not regulate longitudinal groundspeed (“unregulated groundspeed”, right column). **C**. Model C, in which each fly regulates longitudinal groundspeed, and maintains a constant trajectory (“fixed trajectory”, middle row) relative to some external azimuthal reference (depicted as a mountain). **D**. Model D, in which each fly has unregulated groundspeed and a fixed trajectory. **E**. Model E, in which each fly controls its heading by means of a random Lévy process (“Lévy flight”, bottom row) and regulates its longitudinal groundspeed during each straight-line segment of the Lévy flight. For graphical clarity, airspeed, trajectory, and longitudinal groundspeed vectors only shown for the second of the three segments depicted. **F**. Model F, in which each fly has unregulated longitudinal groundspeed and adopts headings governed by a Lévy flight.

**Figure 6.**
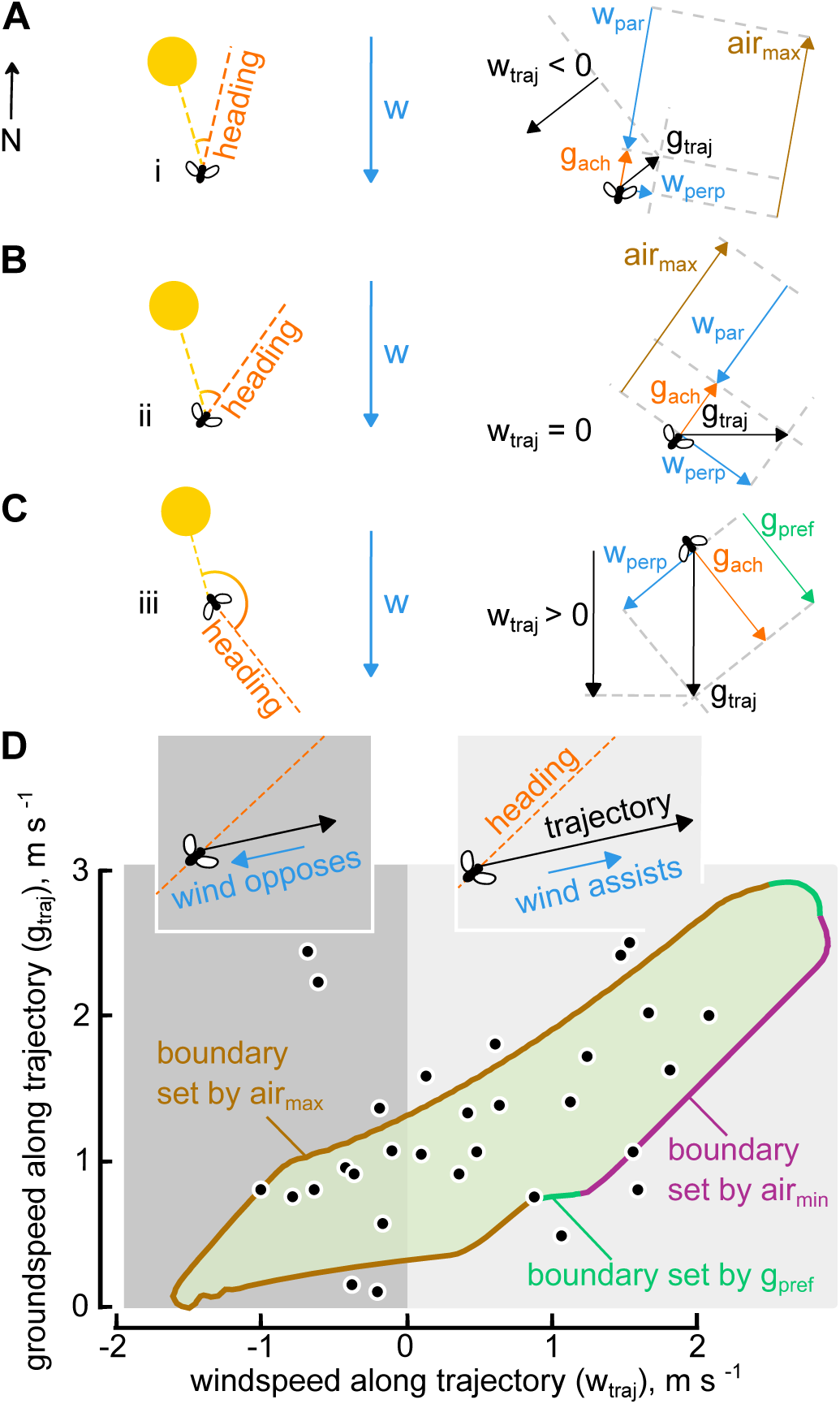
Field data support a behavioral model integrating groundspeed regulation and heading fixation. **A - C**. How the rules of model A, applied to three flies differing only in their heading angle with respect to some azimuthal cue (e.g. the sun), might generate trajectories i – iii depicted in Fig. 4A. **A**. Because fly i holds its heading (broken orange arrow, heading) nearly due north (N), a large component of the wind (w, blue vector) is antiparallel to its heading (blue vector, w_par_). Exerting maximum airspeed (brown vector, air_max_) against w_par_, the fly achieves a modest groundspeed along its body axis (orange vector, g_ach_). This vector sum of g_ach_, with the perpendicular component of the wind (blue vector, w_perp_), yields the fly’s total velocity relative to the ground (black, g_traj)_, the magnitude of which is one of the field-measurable parameters (Fig. 4B, plotted along ordinate). This trajectory is opposed by the wind (black vector, w_traj_), to an extent that is also field-measurable (Fig. 4B, abscissa). **B**. Fly ii orients its body ∼35° from north, and applying the maximum airspeed along this heading results in a trajectory due east. As such, the net trajectory is entirely perpendicular to the wind, yielding w_traj_ = 0. **C**. Fly iii orients its body ∼40° from south, and because in this case w_par_ (vector not shown) is parallel to and of smaller magnitude than the preferred longitudinal groundspeed (green vector, g_pref_), the fly achieves this preferred groundspeed. Summed with w_perp_, this yields a g_traj_ vector pointing due south, and thus a w_traj_ vector of the same magnitude as the wind. **D**. The relationship between the field-measurable parameters described above, w_traj_ and g_traj_, simulated over a range of windspeeds and wind directions, with g_pref_ = 1.0 m s^-1^, air_min_ = -0.2 m s^-1^, and air_max_ = 1.8 m s^-1^. The 99% prediction contour from all simulations is filled in green. Negative values along the abscissa (dark gray) depict simulations for which the fly’s trajectory was opposed by the wind; positive values (light gray) depict trajectories assisted by the wind. Boundaries of this contour are constrained by model parameters air_max_ (brown), air_min_ (purple), and g_pref_ (teal). Black points indicate field measurements from 30 traps over 5 experiments.

To quantitatively compare the performance of the four models, we calculated the average pair-wise log likelihood ratio between model A and each of the alternate models, determined using 40,000 bootstrap iterations (Fig. 7). The resultant distributions of log likelihood ratios indicate that models A and C — both of which invoke groundspeed regulation — predict our field data equally well, and that both match the field data much better than do models B and D, which lack groundspeed regulation. To determine whether these conclusions were robust, we performed a sensitivity analysis in which we ran simulations across a wide range of values for each of the free parameters (Supplemental Figure 3). For models A and C, having three free parameters, we ran 864 simulations; for models B and D, with one free parameter, we ran 10 simulations. From this, we determined the parameter set for each model that best fit the field data. We then performed pairwise comparisons of the individually optimized models, using the same metric described above (Fig. 7). As before, we found that the two models invoking groundspeed regulation better explained the field data than the models lacking this feature (Supplemental Figure 4). The optimized parameter values were quite similar to the ones we had chosen *a priori* (as in Fig. 7) based on estimates from the literature; model A was optimized by air_max_ = 2.0 m s^-1^, air_min_ = -0.5 m s^-1^, and g_pref_ 1.25 m s^-1^; model C by air_max_ = 2.0 m s^-1^, air_min_ = -0.2 m s^-1^, and g_pref_ 1.5 m s^-1^. Collectively, our results suggest that a relatively simple behavioral algorithm involving either fixed random heading or trajectory, regulated groundspeed, and unregulated sideslip can account for the salient features of dispersal behavior under a range of different wind conditions. We also created two additional models (D and E, Fig. 5) in which the azimuthal orientation was determined not by either fixed heading or trajectory, but rather according to a Lévy process (26, 27). Both Lévy models predict our field data quite poorly, regardless of whether it does (Fig. 7E) or does not (Fig. 7F) incorporate groundspeed regulation.

**Figure 7.**
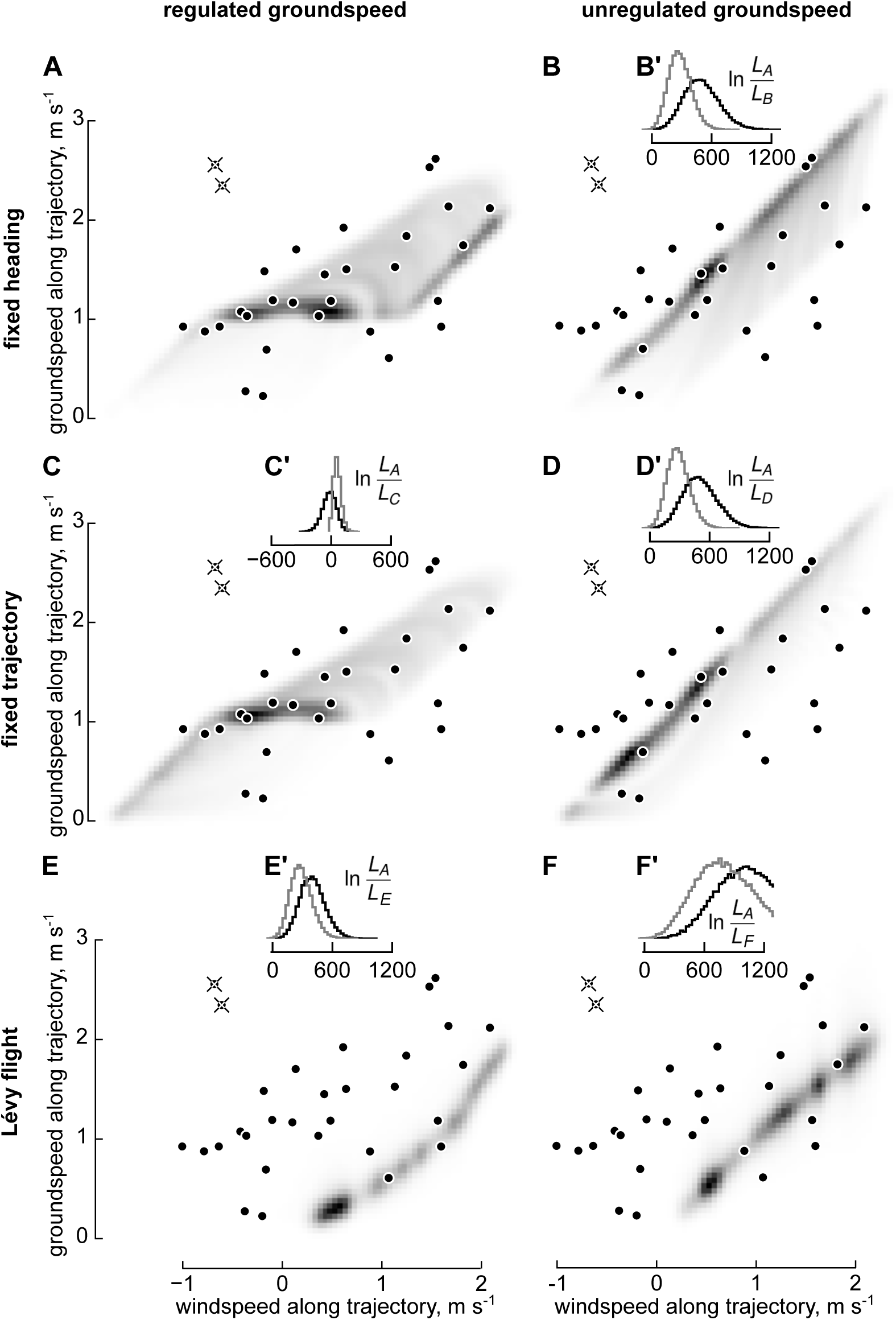
Behavioral models incorporating longitudinal groundspeed regulation fit field data better than alternate models lacking groundspeed regulation. **A-F**. Six different behavioral models, simulated over a wide range of wind conditions, generate distinct relationships between two field-measurable parameters, windspeed along average flight trajectory (abscissa) and groundspeed along trajectory (ordinate). Grayscale shading shows each models’ normalized probability density function (PDF), darker denoting higher probabilities. Field measurements from 30 traps over five experiments (closed circles) are plotted over each model’s PDF; likelihood values at each point were compared pairwise between models. In an alternate analysis, two points considered possible outliers (overlaid with crosses) were excluded from the likelihood-ratio calculation. **A**. PDF generated by model A, in which each fly maintains a fixed body angle relative to an external azimuthal reference, and adjusts its airspeed (within min and max limits) to achieve a fixed groundspeed along its body axis. **B**. PDF generated by model B, in which each fly maintains a fixed heading as in the previous model, but does not regulate groundspeed along its body axis. **B’**. Bootstrapping the field data over 40,000 iterations generated a distribution of log likelihood ratios comparing model A to this model. Positive values denote iterations in which model A predicted the resampled data better than did model B. The mean of this distribution was 499 (black histogram); when re-calculated excluding the two outlying data points, the distribution had a mean of 282 (gray histogram). **C**. PDF of model C, in which each fly has a regulated longitudinal groundspeed, and maintains a constant trajectory relative to an external azimuthal reference. **C’**. As in B’, but comparing models A and C. Distribution mean, -20; excluding outliers, 61. **D**. PDF from model D, in which each fly has unregulated groundspeed and a fixed trajectory. **D’**. As in B’, but here comparing models A and D. Distribution mean, 504; excluding outliers, 288. **E**. PDF of model E, in which each fly regulates longitudinal groundspeed and controls its heading via a Lévy-flight foraging algorithm. **E’**. Here comparing models A and E. Distribution mean, 400; excluding outliers, 287. **F**. PDF of model F, wherein each fly adopts headings via the same Lévy-flight algorithm as the previous model, but does not regulate its groundspeed. **F’**. Here comparing models A with F. Distribution mean, 1076; excluding outliers, 828.

In addition to evaluating our six models with respect to trap-arrival dynamics, we also tested their ability to predict the overall angular distribution of trap counts with respect to the wind (Fig. 2C). Our simulations provide angles of each fly’s trajectory, so we examined the circular variance of these as a function of windspeed for each of the six models. All six models show a narrowing of the population trajectories as wind increases (Supplemental Figure 2), qualitatively consistent with our field data. However, because we made no attempt to model the process of plume tracking – which we expect to strongly affect final trap counts in the field – it is beyond the scope of the present study to use final trap-count distributions as a metric for quantitatively comparing different models’ fit to our field data.

## Discussion

We conducted a series of release-and-recapture experiments to quantify the dispersal performance of *Drosophila melanogaster* under field conditions. Our results are consistent with the prior results of Coyne and co-workers(8), who concluded that *Drosophila* could disperse up to 15 km over a single night. Using a circular array of traps equipped with cameras, were able to examine the influence of wind on the overall movement pattern of the population, and found that whereas flies tended to fan out evenly in all compass directions at low windspeeds, their flights were biased more downwind as wind strength increased. Collectively, the data from all of our releases were consistent with a single behavioral algorithm in which each fly chooses at random and maintains a heading (or trajectory), regulates groundspeed along its longitudinal body axis, but tolerates wind-induced sideslip.

We deliberately chose to conduct our experiments on a flat, featureless, dry lake bed to simplify the external factors that might influence dispersal behavior, such as visual and olfactory signals from nearby vegetation. However, we acknowledge that our experiments were artificial in a number of important ways. First, *Drosophila melanogaster* is a cosmopolitan species that originated in Africa(28) and is not native to the Mojave Desert. Thus, we can infer little regarding the species-specific behavior of the flies as it relates to their ancestral habitat, or habitats that the species has more recently invaded. Instead, we suggest that our observations more likely shed light on deeply rooted behavioral algorithms that are shared by many insects and not the result of recent evolutionary processes(29).

Second, like all release-and-recapture protocols, our experiments forced a rapid, mass exodus, whereas dispersal within the normal life history of flies is more likely a choice influenced by a complex interaction of internal and external factors. In some insect species, dispersal is a distinct behavioral syndrome accompanied by morphological specializations, such as increased wing length, that are triggered during ontogeny(30). Even in species in which dispersal is not associated with distinct morphological changes, it may be anticipated by physiological modifications(31). There is no evidence that *Drosophila* exhibit distinct morphological changes associated with a dispersal state, although photoperiod and temperature do alter physiology and body size (32, 33). Further, laboratory populations subjected to strong selection for dispersal ability (as measured in a walking assay) exhibit heritable changes in behavior(34). Thus, it is possible that distinct populations of flies might respond differently in a mass release depending upon their genetic composition. In our experiments with lab-reared individuals, we did not know whether the flies were in a physiological state that would promote dispersal, or whether their genetic background was particularly conducive to dispersal or not. However, upon emergence we maintained the flies *ad libitum* on a protein-deprived diet. This was a deliberate attempt to proffer them an ample energy source while also providing a strong motivation (to the females at least) to search elsewhere for protein-rich food that was suitable for reproduction(35). The fact that the vast majority of flies left the containers upon release gives us some confidence that the animals were not inhibited from initiating long-distance flight by either physiological or genetic factors and that our feeding regime may have had the desired effect.

Our results indicate that dispersing *Drosophila* actively regulate their longitudinal groundspeed to a value of ∼1 m s^-1^ (3.6 km hr^-1^). In his remarkable, ‘ultralong’ tethered-flight experiments(36), Karl Götz demonstrated that fully fed *Drosophila* could fly continuously for up to 3 hours, a value that is consistent with other measurements of metabolic rate and aerodynamic power requirements(37). This suggests that a single fly could cover ∼12 km without the need for refueling, a distance of about 4.8 million body lengths (for comparison, a marathon is about 25 thousand human body lengths). Even if this value represents the extreme of their performance range, it nevertheless demonstrates a large dispersal capacity for such a small insect. There is nothing to suggest that *Drosophila* is particularly unique with respect to its flight biomechanics or physiology, so such performance is probably representative of many species. At the higher groundspeeds we measured under windy conditions (∼2.5 m sec^-1^), the dispersal distance estimate for a single flight increases to 27 km. Flies could almost certainly disperse much further in an uncontrolled fashion in more extreme conditions, especially if they were carried well above the boundary layer into higher elevations where they catch prevailing winds(38). Presumably, it is just such uncontrolled events that resulted in the colonization of isolated oceanic islands by *Drosophila spp*., such as has occurred in the Seychelles and Hawaii(39).

Many animals – ranging from ants(40) to albatross(41) – localize food by moving upwind within advected odor plumes, a behavior termed odor-mediated anemotaxis. This behavior typically involves an iterative sequence of upwind surges when odor is detected, interspersed with crosswind casts when the odor is lost(11). Although we could not directly observe plume tracking in our field experiments, *Drosophila* readily exhibit such behavior in wind tunnels(17, 42). Whereas the utility of the cast-and-surge algorithm is obvious once an animal encounters a plume, the best flight path an animal should choose with respect to the wind before it detects an odor is less clear, and may depend on wind conditions(43, 44). Some animals, such as the cabbage root fly, appear to fly upwind when experiencing an odorless background flow(45). This might be logical, simply because if an animal detects an odor, its source must be upwind. On the other hand, by flying upwind an animal is limiting itself to odor targets that reside in a narrow sector directly ahead. The propensity for upwind or downwind orientation when flying in wind tunnels should be viewed with caution, because the tunnel geometry prohibits extended motion in all but these two directions(42). Tsetse flies fly do appear to fly downwind prior to detecting an odor plume(46), possibly because this allows them to cover a greater distance while searching for hosts. Perhaps the most logical search strategy is to deliberately move crosswind, because such a trajectory would potentially intercept the largest number of upwind plumes. Such a strategy has been observed in both desert ants(40) and soaring albatross(41).

In our experiments, we observed that flies fan out in all directions at low windspeeds, similar to the behavior reported in gypsy moths(47), while at higher speeds they are biased downwind. The behavior we observed, and the model we propose, may be viewed as a compromise between the need to find an attractive odor plume and the goal of using the wind to increase dispersal distance. By not regulating sideslip, an insect allows itself to be directed downwind, but by regulating forward groundspeed, it maintains some crosswind component that might increase the probability of encountering an upwind plume. As windspeed increases, the behavior converges on pure advection, but under such conditions it less likely that the animals would have the capacity to track a plume upwind even if they detected one, at least until the windspeed decays.

Although we captured flies at odor-baited traps, there is some possibility that the flies arrived without having tracked its associated plume upwind. If 80,000 flies fanned out evenly in all directions from the release site, they would reach the perimeter of the trap radius at a linear density of ∼12 flies per meter. If we liberally assume that a fly might be able to see a trap from a distance of 10 meters, then it is possible that ∼240 flies would pass near enough to a trap that they might land on it without needing to follow the odor plume. Thus, we may have only trapped flies that happened to choose trajectories that carried them near one of the traps. There are several arguments against this interpretation. First, our coarse estimates for how many flies came within the visual detection range of the traps are almost certainly an overestimate as it assumes that all the flies flew within 1 to 2 meters of the ground where they could encounter the trap. *Drosophila spp*. have been observed at high density in 200-m high aerial traps(39) and it is possible that many of the flies in our experiments rose well above the ground after release. Second, laboratory experiments indicate that flying flies are not attracted to land on visual objects until after they have encountered an attractive odor(17), thus seems reasonable to assume that the flies would have made some contact with the plume before landing on the trap.

However, our results do suggest that the distance at which a fly can successfully detect or track an odor plume is limited. If flies could easily detect and track plumes over distances of 1 km or more, then we would have expected to see a recapture bias at upwind traps, at least during experiments when wind speeds were low. This is because as the flies radiate from the release site, the very first plumes they are likely to contact originate from upwind traps. The fact that we could not detect an upwind bias implies that the maximum detection distance of the plumes we generated must be substantially lower than 1 km, perhaps on the order of 100 m or less. Accurately determining the length scale at which flies can detect and track a plume is a high priority of our future studies, but will require a substantially different experimental design.

Of the models we tested, the two that combined longitudinal groundspeed regulation with either fixed heading or fixed trajectory (panels A and C in Figs. 5 and 7) performed best in predicting our field data. Of these two, we believe that the fixed heading model is the most biologically plausible. Our result that flies tend to fan out in all directions at low windspeeds is consistent with recent laboratory experiments showing that flies adopt arbitrary headings relative to patterns of skylight polarization and sun position. Although tethered *Drosophila* will also steer toward large conspicuous visual objects (30) – a reflex called stripe fixation – it is unlikely that this reflex could explain the behavior of the flies on the lake bed. First, although the lake bed is surrounded by some ridges (Fig. 1A), none contain vertical features that seem prominent enough to elicit the fixation response required to generate a constant trajectory. Even if there were one or two geographic features large enough to attract the flies, this could not easily explain how flies fanned out in all directions. As argued elsewhere, stripe fixation is better interpreted as a transient attraction toward nearby objects rather than a long distance orientation behavior(29).

Dispersing *Drosophila* use celestial cues not as a compass to go in a preferred direction, but rather simply as a means of holding an arbitrary orientation (i.e. menotaxis). In this regard, flies’ use of the sky compass system is similar to dung beetles, which maintain a straight trajectory away from the dung pile(48), but not like monarch butterflies, which use the sun compass to fly in a particular direction(49). However, as has been pointed out by Honkanen and coauthors (50), the alteration in central complex circuitry that would be required to transform compass-based random dispersal behavior into seasonal migration might be quite subtle, and the two behaviors are perhaps better viewed as points on a continuum. This notion is supported by field measurements of the navigational accuracy of monarch butterflies; even these virtuosic navigators fan out quite widely during their migration(51), albeit with seasonally appropriate geographic bias.

In our experiments, the released flies were deprived of any attractive sensory cues until they chanced upon an odor plume originating from the traps. Thus, the early stages of our experiments closely match the critical assumptions that underlie the theoretical search behavior known as Lévy flights. According to this theory, a searching agent deprived of any sensory cue emanating from its targets should implement a sequence of randomly directed straight flights, the length distribution of which exhibit power law statistics (22). The free flight trajectories of *Drosophila* flying within an enclosed chamber have been interpreted to support Lévy flight theory(52),

although a subsequent study argued that flight segments within such chambers are truncated by vision-based collision avoidance reflexes triggered by the looming wall, and thus the resulting run distribution cannot be interpreted as resulting from random events(53). Our simulations that incorporated a Lévy process (panels E and F in Figs. 5 and 7) yielded the worst match with our field data, largely because flies executing the zig-zagging flight path take much too much time to cover the 1-km distance from release site to trap. The parameters we chose in the Lévy simulation were generous, in that the minimum and maximum run durations were 1 s and 1000 s, respectively, which are substantially right-shifted (i.e. favoring long runs), compared to measured rates of spontaneous saccades(54). Thus, our results provide strong evidence that flies do not employ a Lévy process during dispersal, even though the conditions of our experiments almost perfectly implement the underlying assumptions of the theory.

In summary, we propose a model of wind-assisted dispersal in which each insect chooses and maintains a random heading, regulates its longitudinal groundspeed, but tolerates wind-induced sideslip. While undoubtedly simplistic, the advantage of our model is that it can explain dispersal behavior under a variety of wind conditions without requiring that any individual animal change its behavioral setpoint as a function of windspeed. It thus represents a biologically feasible ‘rule-of-thumb’ that yields a desired behavioral outcome without requiring sophisticated neural computations. Although derived from measurements on *Drosophila*, we suggest that the model might explain the dispersal behavior of many flying insects with roughly similar natural histories.

## Materials and methods

### Release chamber and flies

We fashioned the release chambers from 66 × 41 x 34 cm plastic containers (Sterilite, 1757), which we modified in several ways. To allow us to insert our arms into the chambers without releasing flies, we cut a circular hole in the side of one wall, and fitted it with a long cylindrical cloth sleeve. We drilled small holes into the container lid to permit the flow of air, and lined the inner surface of the lid with fabric, which prevented escapes. This lining consisted of two layers, the inner a polyester mesh (Joann Fabric, 400075440594), and the outer, a tight-woven cotton less permissive to airflow, that we cut into adjustable flaps. To provide food to the adults during storage and transport, we poured a 1-cm thick layer of sucrose agar (16 g/L sucrose, 6.8 g/L agar, 0.75 g/L CaCl_2_, 0.73% EtOH) in the bottom of the chamber, which provided an abundance of water and carbohydrates, but very little protein.

The flies we used in this study were descendants of 200 wild-caught *Drosophila melanogaster* females maintained in our laboratory since 2013. To expand the population for each release chamber prior to each release, we placed approximately 10 gravid females in 100 - 140 plastic fly bottles (225 ml) filled with standard cornmeal medium. To transfer flies to the release chambers, we lined the inside of each of the fly bottles with transparency film (3M, CG6000) that we laser cut to closely conform to the inner surface of the bottles, above the level of the food, when placed inside (Supplemental Figure 5A, B). During wandering stage, the larvae would crawl upward and pupate on the inner surface of the sheets, after which we removed these plastic inserts laden with pupae and hung them like coat hangers on horizontal tubing strung in the release chamber for this purpose. We did not quantify post-pupal mortality rates, but we estimate that 80 – 95% of transferred pupae remained healthy through the day of the release.

After transferring the pupae, we maintained the release chambers at room temperature, but sometimes placed the chambers in an incubator set to adjust flies’ developmental timing relative to permissive weather forecasts at the field site. We regulated humidity within the chambers by visually checking for condensation on the chamber walls and accordingly opening or closing the outermost cotton flaps. Prior to transporting the flies to the field site, we removed the plastic transfer sheets and hung fabric strips from the horizontal tubing to provide the flies with ample surface on which to perch. During transport in an air-conditioned car, we covered each chamber with a reflective tarp to reduce heating.

### Field site and trial protocol

We conducted the release-and-recapture experiments at Coyote Lake, a dry lake bed in the Mojave Desert (Fig. 1A), after receiving a permit from the United States Department of the Interior, Bureau of Land Management, Barstow Field Office. We chose this location because the lack of vegetation within the playa provided a relatively simple visual and olfactory environment. For most experiments, we deployed ten camera traps at a 1-kilometer radius from our fly release site located at 35.05883° latitude, -116.74556 ° longitude. In one initial experiment, we deployed 8 traps at a 250 m radius.

The traps were fashioned by modifying 28 × 28 x 17 cm plastic containers with a detachable lids (Container Store, 10062800). The surface of each trap consisted of a 28 × 28 cm white polyester woven mesh surface (Joann Fabric, 400075440594) interdigitated with a hexagonal array of 60 mesh funnels projecting downward into the trap cavity. Each funnel entrance had an oval shape of 12 × 18 mm and extended 15 mm into the trap, terminating with an aperture of 2.5 mm. We cut the fabric pieces for the funnels using a laser cutter, sealed the seams with the laser cutter, and then sewed them onto the mesh surface of the trap. When deployed in the field, the mesh top was pulled taut over the top of the container and secured by attaching the original plastic lid in which everything but the sealable rim had been removed.

We baited the traps with a 500 mL solution of apple juice (Tree Top), champagne yeast (Lalvin EC-1118, 1 gram L^-1^) and granulated cane sugar (Domino or C&H brand, 154 grams L^-1^). The mixture was allowed to ferment at 24°C for 1.5 to 2.0 days, and on one occasion for 3.0 days, at which point it provided a source of ethanol, CO_2_, and other attractive odors (van Breugel *et al*. 2018). The variability in fermentation time was due to the fact that we could not perfectly predict the wind conditions at the field site, and occasionally had to conduct a release later than originally planned. We poured the ferment into the traps as we deployed them in the field, and also added a small tub of banana puree (Gerber, 113 grams) to provide flies that entered the traps with an attractive place to land.

For each experiment, we positioned the release site and the surrounding ring of traps using a handheld GPS device with stationary accuracy of 2.5 m. We set up an anemometer (MetOne, direction sensor 020C, speed sensor 010C) at the release site, at a height of ∼2 m above the ground. We oriented the direction sensor azimuthally using a magnetic compass and later adjusted for declination. We logged anemometer data at 20 Hz on a Raspberry pi computer with custom-written software.

After deploying the anemometer and traps, we drove our two vehicles approximately 1.5 km away from the release site to limit visual and olfactory stimuli within the experimental arena, leaving one or two people to release the flies. In our initial experiments, we released one chamber of flies; in later experiments, we generally released two chambers at the same time. Five minutes after releasing the flies, we sealed the chamber(s) in plastic bags in an effort to contain olfactory cues and the remaining flies. Approximately one hour after the release, we began collecting the flies in the traps (Fig. 1C), which we preserved in ethanol. After transporting the preserved flies to the laboratory, we counted the number of *D. melanogaster* and inspected the collection for local species that might be confused with *D. melanogaster* in our camera images. The only other drosophilid fly we found in our traps over the course of the all experiments, was *Drosophila suzukii*, of which we collected a total of 7 individuals over all the releases.

We monitored our baited traps with cameras (Raspberry pi, Pi NoIR Camera V2) mounted 27 cm above each trap, which took time-lapse images of the trap surface at 0.5 or (in some experiments) 0.33 Hz. We controlled the cameras with Raspberry pi computers, on which we had installed an interface (https://github.com/silvanmelchior/RPi_Cam_Web_Interface) to control camera parameters. Among all cameras and our anemometer logger, we maintained clock synchronization within ∼1 second by outfitting each Raspberry pi with a real-time clock (Adafruit, DS3231) (Fig. 1C).

## Data analysis

### Estimating size of release population

To estimate numbers of flies released, we counted the pupae on a subset (∼30%) of the pupal transfer sheets by taking digital images and subjecting them to a simple machine-vision analysis using OpenCV (3.3.1) for Python. We generated a binary image via adaptive thresholding (threshold and neighborhood area determined empirically, against manual annotation), from which we detected contours. Each image yielded a histogram of contour sizes, and we used the position of the prominent, small-pixel-area peak to estimate the size of a single pupa. Dividing the total contour area by the single pupa area provided our estimate of the number of pupae on that sheet.

### Quantifying fly arrivals at baited camera traps

We analyzed the time lapse images from each camera with custom-written machine-vision software, supplemented by human annotation. The first step in our machine-vision pipeline was applying a binary mask to exclude the lakebed surface and any corners of trap fabric that might move in the wind. We then ran the masked color image stack through a mixed-Gaussian background subtractor (OpenCV 3.3.1), trained on a sliding window (typically 25 frames) to account for moving shadows. To avoid false negatives in cases in which flies paused on the trap surface, we interposed one or two frames between this background-training window and the analysis frame from which foreground objects would be detected. We detected foreground pixels on the basis of their squared Mahalanobis distance (typically 10 - 25) from the background model. We excluded brighter-than-background pixels and then smoothed and detected contours from this binary image. We classified each contour as: (1) a fly atop the trap, (2) a fly within the trap, or a (3) non-fly on the basis of its area, eccentricity, and contrast relative to surrounding pixels.

For each trap analyzed, we used a graphical interface to manually evaluate frames annotated by our automated analysis. We generally found good agreement for the scoring of flies atop the trap, but higher rates of both false positives and false negatives for flies within the trap, which is not surprising given that such flies were obscured beneath the mesh surface. Our automated analysis excluded insects that were either too small or too big, but was generally unable to exclude *Drosophila*-sized insects (e.g. a small beetle) even if a human viewer could easily identify them from the digital image as a different species. Such misclassifications often led to a background level of false positives, essentially a noise floor on which the arrival wave of released flies was superimposed. To better estimate the start of the arrival wave, we smoothed the on-trap data with a sliding mean of 10 frames, and calculated the mean and standard deviation during the first 200 seconds after the release of the flies, which we considered as the characteristics of our noise floor immediately prior to the arrival of the flies. We then used these statistical values to set minimum and maximum values (0.2*(mean + 2.0 STD) and mean + 2.0 STD, respectively) for a Schmitt classifier. This classifier binarized our time-series data into bouts above, or below, the noise floor. The start of the longest bout above the noise floor was declared the time of the arrival wave. In addition to estimating the start of the arrival wave with our automated analysis, we manually scored the first arrival of a *Drosophila-*shaped insect in the image stream from each trap, starting at 200 seconds after the fly release and continuing until we unambiguously detected the first fly (Supplemental Figure 1). While manually scoring, we occluded the time stamps on each image to reduce observer bias.

For each trap, we used the arrival time of the first fly to calculate average groundspeed along (*g*_*traj*_), by assuming a minimum travel distance of 1 km. The average windspeed along the trajectory (*w*_*traj*_) was calculated as:

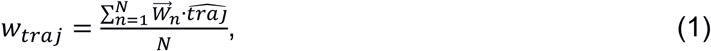

where 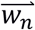 is the wind velocity vector and 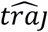 is the unit vector of the average trajectory to the trap. The summation includes all the wind measurements made from the time of release (n=1) to the arrival time of the first fly (n=N).

### Simulations of six behavioral models

Each simulated run of our dispersal models consisted of calculating the resultant vectors according to the rules governing the flies’ azimuthal orientation (Fig. 4). For models A and B, we fixed the flies’ heading; for models C and D, we fixed the flies’ trajectory; for models E and F, we dynamically updated the flies’ heading according to a Lévy flight algorithm. To generate the variation in wind that mimicked the field conditions, we permutated the direction and speed of the simulated wind in each run, using 180 wind directions linearly spaced around a circle, and 261 windspeeds, sampled randomly from a probability density function measured in the field. This probability function was generated non-parametrically using a kernel density estimate (Gaussian kernel with a standard deviation of 0.1 m s^-1^) of vector-averaged windspeeds, w_ave_, calculated from each of the 30 field data points used to evaluate the models:

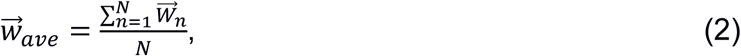

in which the summation interval includes all wind data points from the time of release (n=1) to the arrival time of the first fly (n=N).

Our models imposed maximum and minimum limits on the airspeeds that flies could actively exert along their body axis (air_max_ = 1.8 m s^-1^, air_min_ = -0.2 m s ^-1^). In the case of models A, C, and E, flies regulated the groundspeed along their body axis to a preferred speed of 1.0 m s^-1^, and in models B, D, and F, flies simply exerted a forward airspeed of 1.0 m s^-1^. Only under one condition did we allow flies to adjust their airspeed in models B, D, and F: if the wind was blowing them backwards along their body axis, in these cases we allowed flies to increase airspeed up to airma_x_. We assumed that flies did not tolerate negative groundspeeds parallel to their body axis (i.e. they would not allow themselves to fly backwards); in all models, if backwards flight was unavoidable given the set airspeed limits, we simply dropped the fly out of the simulation. However, we repeated all simulations without this “dropout” assumption and found the that results were qualitatively similar.

For Lévy flight simulations (models E and F), the run-length intervals were randomly drawn from a probability distribution given by:

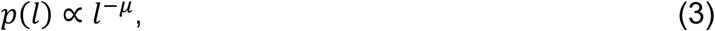

where *l* is run length and *μ* was set to 2.0, a value that is predicted to generate optimal search under many conditions(55), and is close to a value reported from lab experiments using *Drosophila*(52). Between each run, the fly executed an azimuthal turn, drawn randomly from the sum of two Gaussian distributions with means of -93° and +93° and standard deviations of 27°, values based on measurements of spontaneous turns in free-flying *Drosophila*(56).

### Statistical comparisons of six behavioral models

From all 180 × 261 runs of each model, we generated the two-dimensional probability density function (PDF) as a kernel density estimate (σ = 0.05 m s^-1^). The Gaussian kernels were not truncated, to ensure that all field data points overlaid non-zero values of the PDF. We then performed a likelihood ratio analysis, testing whether the five alternate models (B - F) could explain the field data better than model A. The log Bayes factor we report for each pairwise test was calculated as:

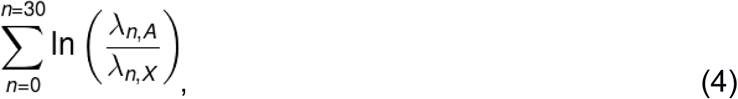

where λ_n,X_ is the likelihood score for data point *n* given its corresponding probability in the PDF of model *X*. We performed this procedure over 40,000 bootstrap iterations of the field data, generating a distribution of log Bayes factors for each pairwise model test. In these distributions, any values below zero indicate iterations in which the alternate model explains the resampled field data better than does model A. We performed these analyses twice, once including and once excluding two suspected outlying field data points (annotated with crosses, Fig 7).

### Sensitivity analysis of our model comparisons

To determine whether our model comparisons were robust to our selection of free parameters for each model, we ran simulations over a wide range of values for each of the free parameters (Supplemental Figure 3). For models A and C, we ran 864 simulations, testing all permutations of 14 values of g_pref_, 14 values of air_max_, and 6 values of air_min_, omitting 312 parameter sets in which g_pref_ would have exceeded air_max_. For models B and D, with one free parameter, we tested 10 values of fixed airspeed. We ran a within-model procedure to determine the parameter set that would allow each model to best fit our data. To do this, we chose one set of parameter values as the reference simulation for pairwise comparisons to each other simulation of that same model; these comparisons assessed model fit to our field data (excluding the two outlying data points) using the mean of a bootstrapped distribution of log Bayes factors, as in Figure 7. After optimizing each model’s parameter values in this manner, we performed pairwise comparisons across the individually-optimized models (Supplemental Figure 4), again using the same metric as described for Figure 7.

### Other summary statistics

To describe the effect of wind on the distribution of final trap counts (Fig. 2), we used both a parametric and non-parametric statistical summary. For the former, we fit a von Mises distribution to the trap counts and report the reciprocal measure of circular dispersion, κ. For the latter, we calculated the circular variance of trap counts by treating each fly as a vector of length one, pointing in the direction of the trap at which it was caught. These vectors were summed and the resultant length was divided by the total number of flies, yielding the vector strength for that release. The circular variance is equal to the vector strength subtracted from 1.

## Acknowledgements

We wish to thank Massimo Vergassola (UC San Diego), who was principal investigator on a grant from the Simons Foundation (71582123) that funded the initial stages of this project. This work was also supported by the NSF (IOS 1547918). Román Corfas, Ainul Huda, Alysha de Souza, Johan Melis, and Aubrey Goldsmith participated in data collection. Annie Rak contributed to preliminary modeling efforts. Bob Verish provided guidance for safely accessing Coyote Lake, and the Barstow Field Office of the Bureau of Land Management permitted our use of this field site.

---

**Supplemental Figure 1.**
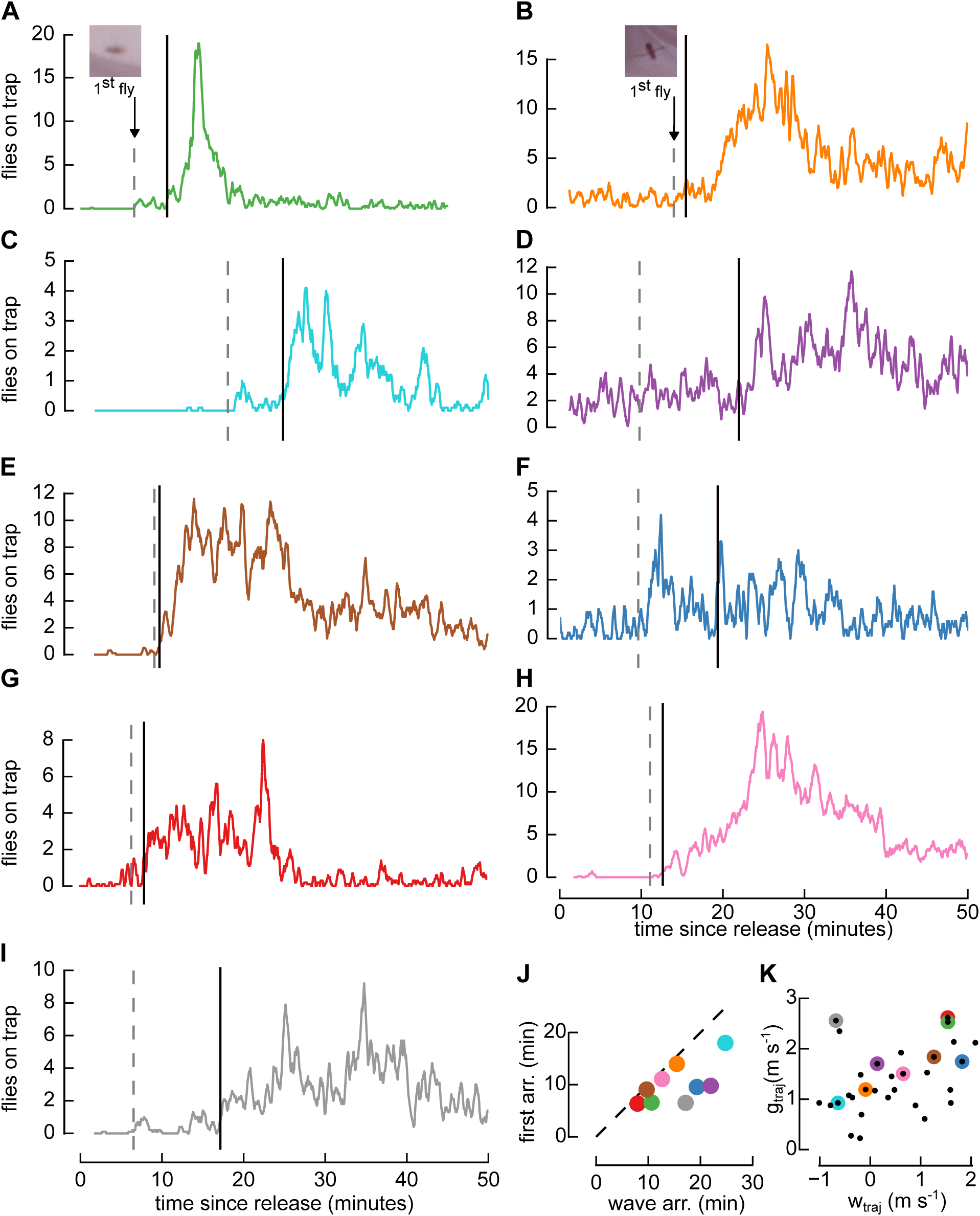
Calculation of arrival time at baited camera traps. **A-I**. Nine examples of trap arrival dynamics scored by machine vision (colored lines), annotated with the time of the first fly arrival as scored by eye (grey broken lines) and with the time of the “arrival wave” (black solid lines) as calculated from the machine vision trace; for A and B, we show the cropped frame of the first fly’s first appearance (insets). **J**. Relationship between the latency to the machine-vision-detected arrival wave (“wave arr.”) and latency to the manually-annotated 1st fly (“first arr.”), overlaid with a unity line (dotted). Point colors correspond to data in A–I above. **K**. The relationship between wtraj (windspeed along trajectory) and gtraj (groundspeed along trajectory) for all 30 data points (black) used previously (e.g. Fig. 7); colored highlights indicate traps whose machine-vision data are presented in A–I.

**Supplemental Figure 2.**
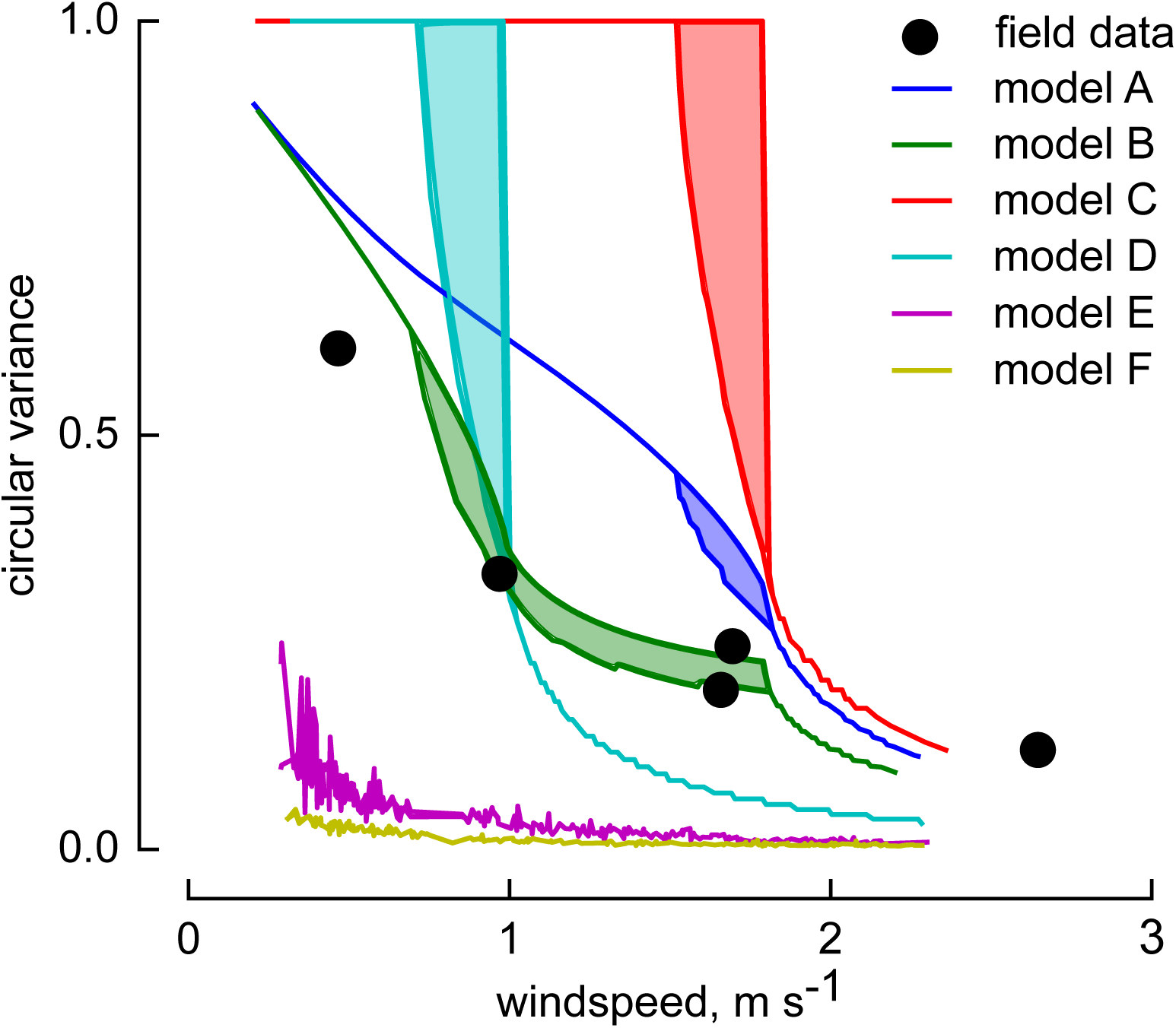
In all behavioral models, the distribution of population trajectories tightens with increasing wind speed. All six behavioral models (solid lines) show a decrease in the circular variance of simulated flies’ ultimate azimuthal trajectories with increasing wind speed. For each model, we calculated circular variance in two ways: first, by simply including all simulated flies (right-shifted), and second, by including only those simulated flies that reached the 1-km trap radius within an hour (left-shifted). The areas between these two lines are filled in for graphical clarity. Our field data (repeated from Fig 2C, black points) are overlaid for reference.

**Supplemental Figure 3.**
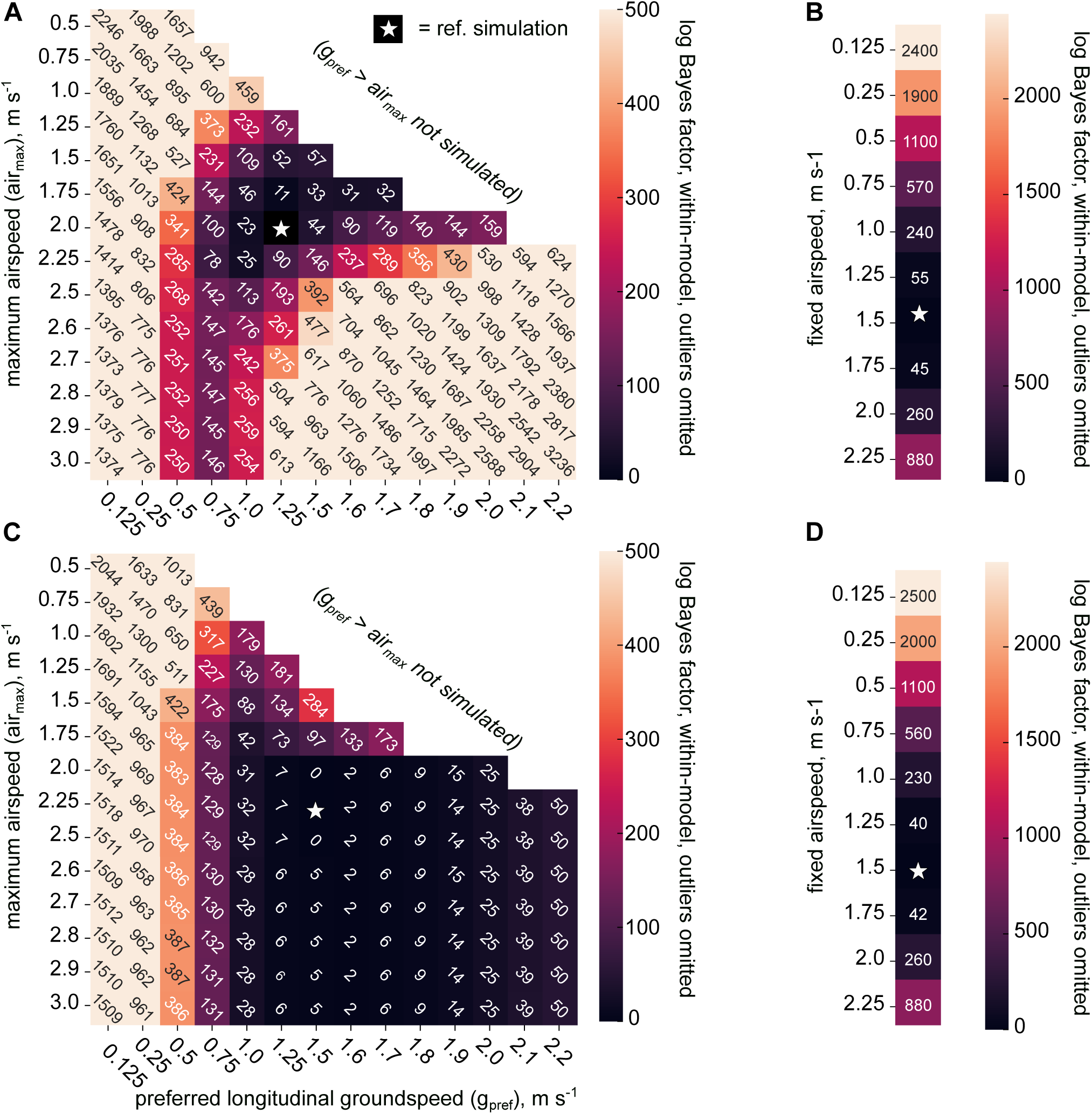
Optimization of each model’s parameters on the basis of model fit to our field data. **A–D**. For each model, one set of parameter values (white star) served as the reference simulation that was compared to each other simulation of that same model, each differing in their parameter values. This comparison examined the relative fit to our field data, using the mean of a bootstrapped distribution of log Bayes factors (color map), just as did the comparisons in Fig. 7. **A**. For model A, varying flies’ preferred groundspeed (g_pref_, columns) and maximum airspeed (air_max_, rows) yields a map of log Bayes factors. The upper-right region, where g_pref_ would be greater than air_max_, was not explored. Lower log Bayes factors (darker colors) indicate parameter pairs allowing the model to perform relatively better in its fit to field data. The third free parameter, the minimum airspeed (air_min_), was far less influential on model fit (data not shown), so we present the map only for a single value of air_min_ = -0.2 m s^-1^. The cell for each parameter pair is annotated with its log Bayes factor. Note that the color map saturates well below the highest log Bayes factors achieved. **B**. Model B, lacking the feature of groundspeed regulation, has only the fly’s fixed airspeed (varying by row) as a free parameter. **C**. Model C has the same free parameters as does model A. Here, also, air_min_ had little effect on model fit, so we show the map corresponding only to the air_min_ = -0.2 m s^-1^. **D**. Model D’s fit to the field data as a function of its single free parameter, the fixed airspeed.

**Supplemental Figure 4.**
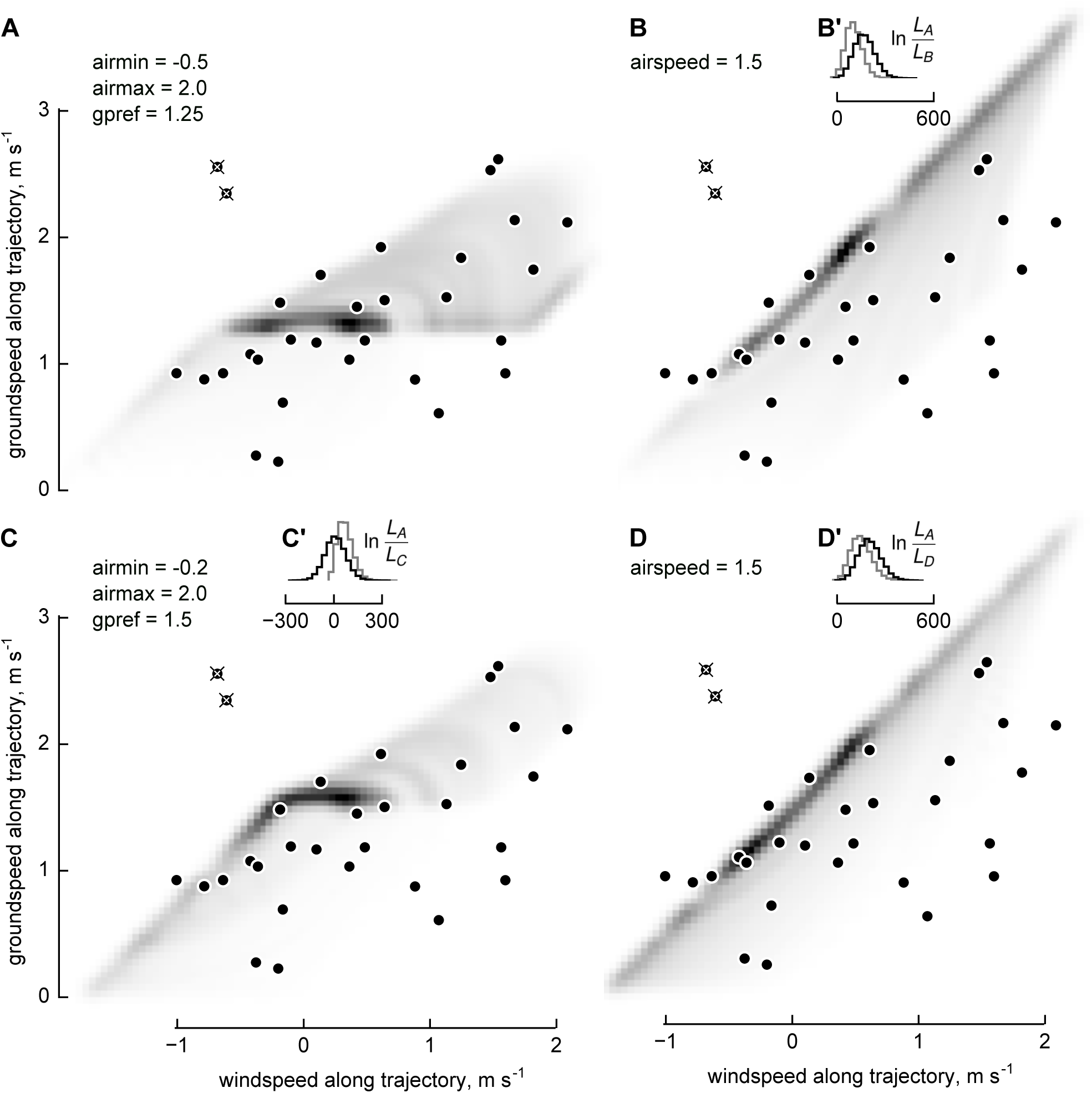
After individually optimizing each behavioral model, cross-model comparisons indicate that models incorporating longitudinal groundspeed regulation continue to best explain the field data. **A–D**. After undergoing parameter optimization (see Supplemental Fig. 3), each behavioral model generates relationships between windspeed along average flight trajectory (w_traj_, abscissa) and groundspeed along trajectory (g_traj_, ordinate). Grayscale shading denotes each models’ normalized probability density function (PDF). Our field measurements (closed circles) are plotted over each model’s PDF; model likelihood values were compared pairwise. In an alternate analysis, two points considered possible outliers (overlaid with crosses) were excluded from the likelihood-ratio calculation. **A**. PDF generated by model A, with its optimized values of air_min_ = -0.5 m s^-1^, air_max_ = 2.0 m s^-1^, and g_pref_ = 1.25 m s^-1^. **B**. PDF generated by model B, with its optimized airspeed of 1.5 m s^-1^. **B’**. Bootstrapping the field data over 40,000 iterations generated a distribution of log likelihood ratios comparing the optimized model A to this model. Positive values denote iterations in which the optimized model A predicted the resampled data better than did this optimized model B. The mean of this distribution was 169 (black histogram); when re-calculated excluding the two outlying data points, the distribution had a smaller variance and a mean of 104 (gray histogram). **C**. PDF generated by model C, with its optimized values of air_min_ = -0.2 m s^-1^, air_max_ = 2.0 m s^-1^, and g_pref_ = 1.5 m s^-1^. **C’**. As in B’, but comparing optimized model A with optimized model C. Distribution mean, 9; excluding outliers, 70. **D**. PDF generated by model D, with its optimized airspeed of 1.5 m s^-1^. **D’**. As in B’, but here comparing optimized model A with optimized model D. Distribution mean, 203; excluding outliers, 150.

**Supplemental Figure 5.**
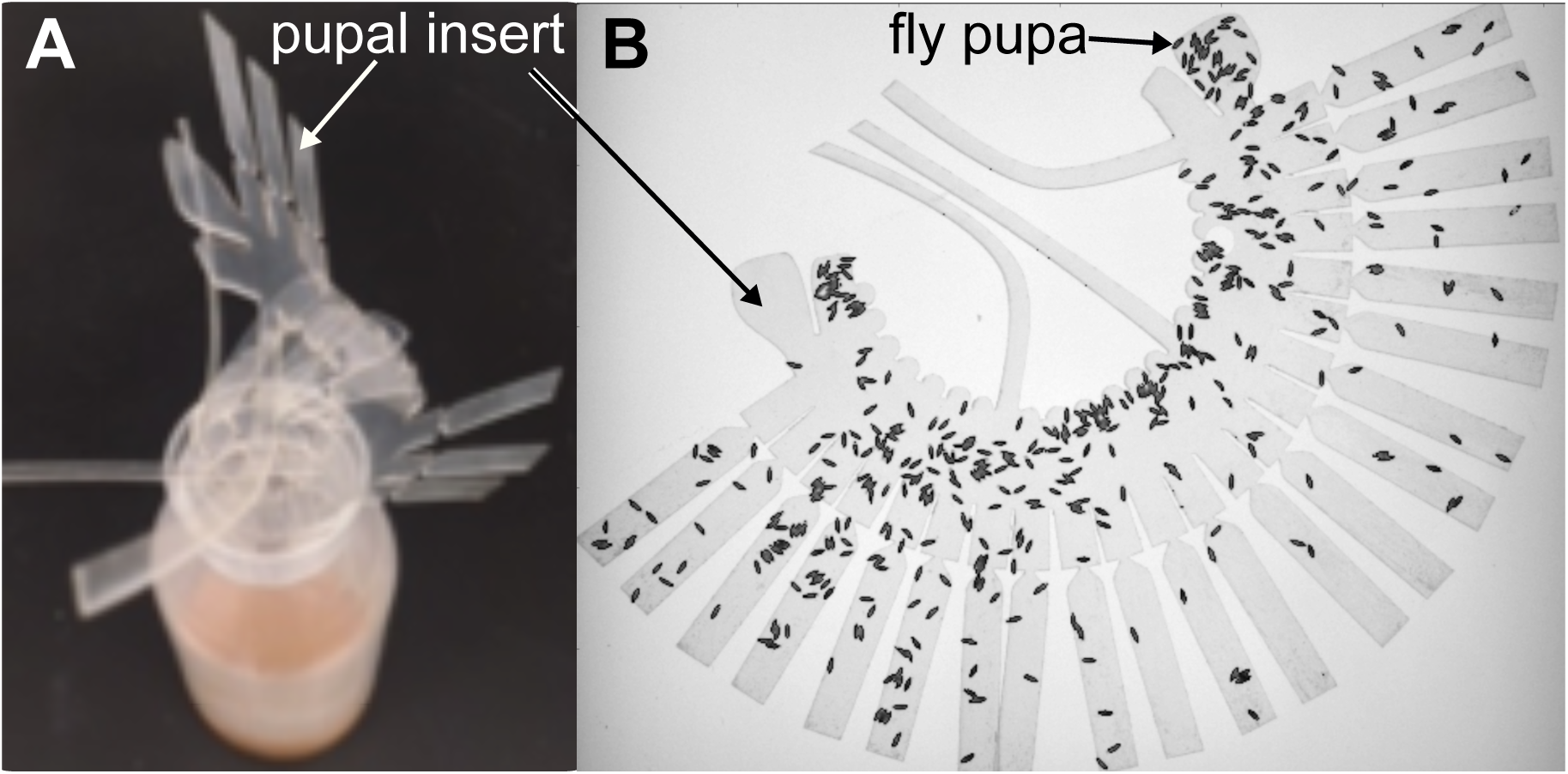
Rearing *Drosophila* for synchronized field release. **A**. Oblique view of a fly-rearing bottle with a pupal sheet partially inserted; when fully inserted, the sheet conforms to the bottle’s inner wall. **B**. A sheet bearing hundreds of pupae, removed from the bottle for imaging and transfer to the release chamber.

